# The unexpected loss of the “hunger hormone” ghrelin in true passerines: A game changer in migration physiology

**DOI:** 10.1101/2023.05.23.541918

**Authors:** Stefan Prost, Jean P. Elbers, Julia Slezacek, Silvia Fuselli, Steve Smith, Leonida Fusani

## Abstract

Migratory birds must accumulate large amounts of fat prior to migration to sustain long flights. In passerines, the small body size limits the amount of energy stores that can be transported and therefore birds undergo cycles of extreme fattening and rapid exhaustion of reserves. Research on these physiological adaptations was rattled by the discovery that birds have lost the main vertebrate regulator of fat deposition, *leptin*. Recent studies have thus focused on *ghrelin*, known as the “hunger hormone”, a peptide secreted by the gastrointestinal tract to regulate food intake, body mass, and other important functions in vertebrates. Studies on domestic species showed that in birds *ghrelin* has effects opposite to those described in mammals, such as inhibiting instead of promoting food intake. Furthermore, a series of recent studies have shown that *ghrelin* administration influences migratory behaviour in passerine birds, suggesting an important role of this hormone in bird migration. However, using comparative genomic analyses we show that *ghrelin* has been lost in the largest avian taxon Eupasseres, after the basic split from Acanthisitti about 50 million years ago. Eupasserines, also known as True passerines, include all but two of the ca. 10,000 known passerine species. We further found that the *ghrelin* receptor (*growth hormone secretagogue receptor, GHS-R*) is still conserved in passerine birds, as indicated by sites under purifying selection and in line with the effects of *ghrelin* administration. Thus, *ghrelin* adds to a list of hormones highly conserved in vertebrates that have lost their main functions in specific taxa. The maintenance of a functional receptor system, however, suggests that in eupasserine birds another ligand has replaced *ghrelin*, perhaps to bypass the feedback system that would hinder large pre-migratory accumulation of subcutaneous fat.

## Introduction

Twice a year billions of birds migrate between breeding and wintering areas, a journey that is very strenuous and often requires stopover events to replenish fat storages. Migratory birds have the unique ability of storing enormous amount of subcutaneous fat prior and during migration (up to 100% the lean body mass in some species, e.g. Bairlein 1991). In mammals, the amount of fat stored in the adipose tissue is mainly regulated through a feedback system by the hormone *leptin*. However, *leptin* has lost this regulatory function in birds (reviewed in Friedman-Einat and Seroussi 2019). The disruption of the well conserved *leptin* feedback system was suggested to be linked to the need of accumulating large fat storages in a short amount of time to sustain long-distance migration (Friedman-Einat and Seroussi 2019). Therefore, recent studies focused on the gastrointestinal hormone *ghrelin*. Ever since its discovery in 1999 (Kojima et al. 1999), a number of studies revealed important roles of this peptide hormone in the central regulation of food intake, body mass, adiposity, glucose metabolism, sleep, anxiety, and stress (Tschöp et al. 2000, Nakazato et al. 2001, Tolle et al. 2002), among several other functions (see Mueller et al. 2015 for a review). The activation of *ghrelin* via acetylation is catalyzed by an enzyme called *membrane bound O-acyltransferase domain containing 4 (MBOAT4*; Kojima et al. 2016). Ghrelin binds as an agonist to the *growth hormone secretagogue receptor (GHS-R)*. There are several forms of *GHS-R*, however, tetrapods share one single, conserved receptor called *GHS-R1a* (Davenport et al. 2005). *Liver enriched antimicrobial peptide 2 (LEAP2*) was identified as an antagonist to *ghrelin* (Ge et al. 2018).

The functions of *ghrelin* in birds have been investigated mainly in a few domestic species, where it appears to have opposite effects compared to mammals, i.e. it inhibits food consumption and down-regulates the build-up of fat storage (Kaiya et al. 2009). Recent studies showed effects of *ghrelin* on food intake and migratory behavior in passerines (Goymann et al. 2017, Henderson et al. 2018, Lupi et al. 2022). Goymann et al. 2017 and Lupi et al. 2022 found effects of *ghrelin* administration on migratory restlessness and departure from stopover sites in garden warblers (*Sylvia borin*) and yellow-rumped warblers (*Setophaga coronata coronata*), whereas Henderson et al. 2018 showed that injected *ghrelin* reduces food hoarding and mass gain in the coal tit (*Periparus ater*). To gain additional knowledge on the involvement of the *ghrelin* system in migratory behaviour of passerine birds, we conducted comparative genomic analyses. Unexpectedly, we discovered that eupasserines, that is the ‘true passerine’ taxon that comprises all but two of the more than 10,000 passerine species, have lost the gene coding for *ghrelin* although they seem to maintain a functional receptor system. These surprising results indicate that in addition to *leptin*, True passerine birds have lost the second main vertebrate regulator of food intake and body mass. This major evolutionary transition might be functional to sustain the need of rapid, repeated gains of large energy stores in small passerines that typically cannot transport all fuel required for the entire migratory journey.

## Results and Discussion

### Presence and absence of genes upstream of the growth hormone secretagogue receptor in eupasserine birds

As a first step, we blasted the RefSeq protein sequences of *ghrelin, MBOAT4, GHS-R and LEAP2* (from the chicken and the rifleman) against the complete avian NCBI Genomes database (wgs). We were not able to detect *ghrelin* in passerine birds (Passeriformes), except for the rifleman (*Acanthisitta chloris*). The split between Acanthisitti and Eupasseres is the oldest divergence within passerines, dated to around 50 MYA (Prum et al. 2015) or 56 MYA (Selvatti et al. 2015). We found *MBOAT4* to be present in several passerine bird genomes (13-18 species, depending on whether the chicken or the rifleman protein sequence was used). Blast hits ranged from 14% to 100% query coverage and from 28% to 78% identity. The chicken and rifleman *LEAP2* protein sequences matched partially to several passerine bird species. Ranging from 75% sequence overlap with 64.41% match similarity for the common starling (*Sturnus vulgaris*) to 39% sequence overlap with 93.10% match similarity for the Zebra finch (*Taeniopygia guttata*). *GHS-R* was present and complete in all investigated bird genomes. We further examined the presence of these genes specifically in the garden warbler (*Sylvia borin*) genome (GCA_014839755.1), for which *ghrelin* had previously been shown to modify migratory behaviour (Goymann et al. 2017). We were only able to find *LEAP2* partially (37% coverage with a 96.9% similarity) and the complete *GHS-R* (100% coverage with 91.5% similarity). Neither *ghrelin* nor *MBOAT2* showed any blast hits.

We further investigated the presence of these genes in passerine birds using ortholog searches in the NCBI protein database. This resulted in similar findings. We only found annotations for *ghrelin* and *MBOAT4* for passerines in the rifleman. Interestingly, while *ghrelin* showed a similar amino acid (aa) length in the rifleman (114 aa, <u>XP_009080522.1</u>) and in the chicken (116 aa, <u>XP_046782141.1</u>), we found *MBOAT4* to be much shorter in the rifleman (174 aa, <u>XP_009071540.1</u>) than in the chicken (424 aa, <u>NP_001186218.2</u>). However, several non-passerine birds showed similarly short *MBOAT4* annotations, such as the red-throated loon (*Gavia stellata*) with 144 aa (<u>XP_009819505.1</u>) or the Northern fulmar (*Fulmarus glacialis*) with 163 aa protein lengths (<u>XP_009571461.1</u>). We found *LEAP2* orthologs for several passerine birds, ranging in predicted protein length from 75 aa to 119 aa, compared to the 76 aa in the chicken (<u>NP_001001606.1</u>). *GHS-R* in the rifleman (344 aa, <u>XP_009077327.1</u>) showed a similar size to the chicken (347 aa, <u>NP_989725.1</u>). In general, the length of *GHS-R* seemed relatively constant with a length of 352 aa in most passerine birds, except for a few outliers such as the Tibetan ground-tit (*Pseudopodoces humilis*, 301 aa) or the Swainsons thrush (*Catharus ustulatus*, 372 aa).

Though unlikely given the different techniques used to assemble the bird genomes on NCBI, we further investigated whether the general absence of *ghrelin* could be due to missassemblies in the genomes. To do so, we shot-gun sequenced three garden warbler individuals with 30x genome-wide coverage. We then mapped the genomic reads to the chicken *ghrelin, MBOAT4, GHS-R* and *LEAP2* nucleotide sequences. We were not able to find a single read mapping to either *ghrelin, MBOAT4* or *LEAP2*, but the entire coding sequence of *GHS-R* showed a coverage of 11x, 20x and 26x, for the three individuals respectively. *SEC13* and *IRAK2* are the flanking genes of *ghrelin* in the chicken genome. We found both genes present in the garden warbler genome, with the distance between exons of the two genes of a similar length in the chicken (∼13kb) and the garden warbler (∼16kb). Recently, Seim et al. 2015 showed that the insertion of an ERVK retrotransposon in exon 0 lead to the inactivation of *ghrelin* in falcons, which they hypothesize is an adaptation to a predatory lifestyle as it might increase food-seeking behaviour and feeding. We did not find any indication for a repeat insertion in the region between *SEC13* and *IRAK2* in the garden warbler. The presence of the two flanking genes on the same chromosome region and the absence of a partial or inactivated *ghrelin* sequence suggest that *ghrelin* has been lost in eupasserine birds, potentially via some sort of chromosomal rearrangement or pseudogenization.

**Figure 1.**
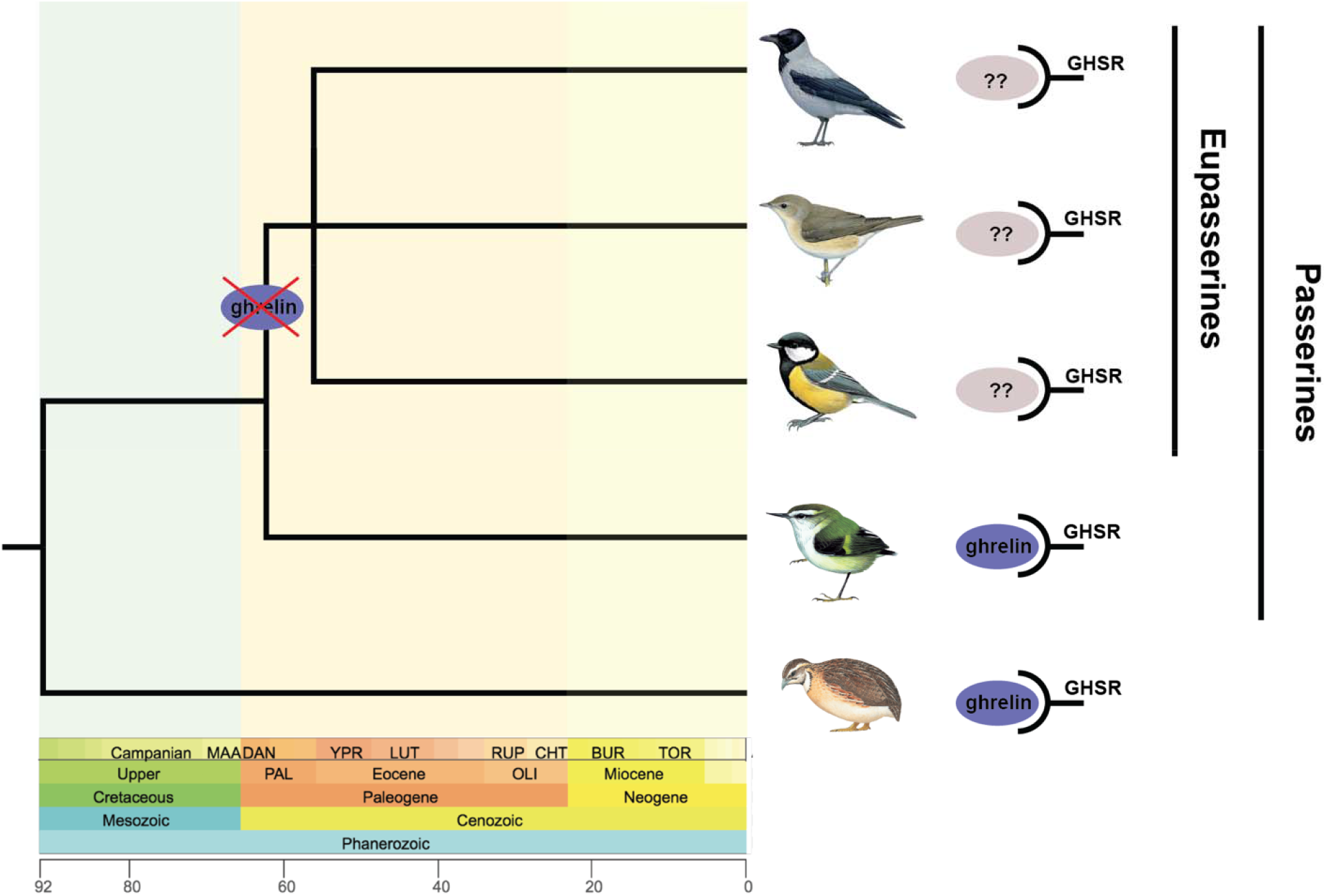
Phylogenetic tree showing the loss of *ghrelin* in eupasserine birds. The phylogenetic tree was obtained from TimeTree (http://timetree.org/; last accessed 21^st^ of April 2023). The x-axis shows the age of the event in million years ago (Mya). Bird drawings were obtained from the Lynx Edicions’ Handbook of the Birds of the World with permission of Alada Gestió Empresarial S.L.

### Evolutionary constraints on the growth hormone secretagogue receptor

Next, we investigated the conservation of *GHS-R* in passerine and non-passerine birds, respectively. The hypothesis would be that *GHS-R* shows little to no conservation in passerine birds if the pathway has been inactivated. We used the two sites selection models FUBAR (Fast, Unconstrained Bayesian AppRoximation; Murrell et al. 2013) andSLAC (Single-Likelihood Ancestor Counting; Kosakovsky, Pond & Frost 2005) to characterize patterns of selection in *GHS-R* using HyPhy (Pond & Muse 2005). Both FUBAR and SLAC indicated high proportions of sites under purifying selection in *GHS-R* (see Table 1), as expected for a conserved coding region. In all cases the SLAC model inferred fewer sites under purifying selection than the FUBAR model. Both methods, however, indicated that passerine birds show considerably fewer sites under purifying selection (FUBAR: 76 out of 357 investigated sites, SLAC: 30 out of 357) than non-passerine birds (FUBAR: 223 out of 357, SLAC: 128 out of 357). To test if this differential amino acid constrain is indicative of a general relaxation of the *GHS-R* gene, we inferred whether the strength of natural selection has been significantly relaxed on the branch leading to the passerine birds, using the RELAX model in HyPhy. The absence of a significant relaxation signature (K=0.89, p=0.784, LR=0.07) and the finding that injected *ghrelin* showed phenotypic effects in passerine birds (e.g. in Goymann et al. 2017, Henderson et al. 2018 and Lupi et al. 2022), indicates that the signaling pathway downstream of *ghrelin* is still functional, and that a different agonist might be regulating this pathway. Preliminary work of our group suggests the existence of a putative ligand with sequence similarities to *ghrelin*-like motifs (data unpublished), which should correspond to the molecule detected in a femtomolar range by radioimmunoassay in previous studies in passerine birds (Goymann et al. 2017; Eikenaar et al. 2018). To date, the nature of the ligand i.e. its affinity to the receptor at the molecular level remains to be determined. However, another possible explanation is that the pathway is a relic and while the receptor is still working, given the absence of the ligand, is on its way to be lost.

### Evolutionary Implications

These and previous findings suggest that in eupasserine birds the signaling pathway including and downstream of *GHS-R* still play an important function in food intake, and thereby maybe even migratory behaviour, but seems to be regulated by a hormone other than *ghrelin*. The structural interactions of *ghrelin* and synthetic *ghrelin* with GSH-R did not change during the evolution of vertebrates, suggesting that *ghrelin* and its receptors evolved separately (Kaiya et al. 2014). This is supported by the finding that to date there is no evidence for multiple *ghrelin* genes in any species (Kaiya et al. 2011).

Previous work has shown that birds diverged from mammals in the way they regulate food intake and fat deposition. The existence and role of *leptin*, the main adipostatic hormone in mammals, in avian species has been debated for decades, until recent studies have shown that in fact birds do not use *leptin* as a feedback signal from the adipose tissue to regulate food intake (reviewed in Friedman-Einat & Seroussi 2019). The authors of the latter paper argue that in birds the ability to fly and migrate might reduce the threat of starvation, therefore unlimited accumulation of fat reserves before migration could be more advantageous for survival than adipostatic control (Friedman-Einat & Seroussi 2019). Eupasserine birds seem therefore to have separated themselves also from the second main system controlling food intake and fat metabolism in vertebrates. Most migratory passerines have a small body size and therefore long-distance migrants cannot carry sufficient energy stores to fuel the entire migratory journey, like several shorebirds do (Battley et al. 2012). Therefore, they rely on alternating rapid fuel deposition at stopover sites and non-stop flights during which they exhaust most of their reserves. This peculiar, repeated alternation between very fat and very lean condition might have required to break away from conserved homeostatic control systems of vertebrates. Gene loss has been reported in a number of taxa as an adaptive mechanism that accompanies major transitions in physiology (reviewed by Monroe at al. 2021), such as for adaptation to aquatic life in cetaceans (Huelsmann et al. 2019; McGowen et al. 2020). Given that migratory behaviour appears to be ancestral to the passerine lineage (Dufour et al. 2020), the release from universal, conserved mechanisms controlling fat deposition and the consequent evolution of novel ways to regulate body mass might be behind the extreme success of this taxon, which alone comprises one third of all avian species and 10% of all tetrapod diversity.

## Materials and Methods

To investigate the presence of *ghrelin* and related genes in passerine birds, we first downloaded the respective protein sequences of *ghrelin* (GenBank accession: NP_001001131.2), *MBOAT4* (GenBank accession: NP_001186218.2), *LEAP2* (GenBank accession: NP_001001606.1) and *GHS-R* (GenBank accession: NP_989725.1) for the chicken (*Gallus gallus*) from NCBI (ncbi.nlm.nih.gov/gene/). We further downloaded the same proteins for the rifleman (*Acanthisitta chloris*): *ghrelin* (GenBank accession: XP_009080522.1), *MBOAT4* (GenBank accession: XP_009071540.1), *LEAP2* (GenBank accession: XP_009082681.1) and *GHS-R* (GenBank accession: XP_009077327.1). We then blasted these sequences against the complete avian NCBI Genomes database (whole-genome shotgun contigs and taxid:8782; last accessed 20^th^ of March 2023) using tblastx for protein sequences (https://blast.ncbi.nlm.nih.gov/Blast.cgi).

Next, we generated genome libraries for three high-coverage garden warbler individuals after a modified protocol following Meyer and Kircher (2010) (detailed laboratory methods can be found in Supplementary File 1). Subsequently, we filtered the raw reads for quality, removed PCR and optical duplicates and trimmed adapters using bbmap v. 38.87 (Bushnell 2014). In order to test whether the absence of *ghrelin* in the passerine genomes on NCBI could be due to assembly issues, we next mapped the genomic reads onto the chicken reference sequences of these genes and *GHS-R* (see above) using BWA 0.7.12-r1039 with the *mem* option (Li and Durbin 2009) and sorted them using samtools 1.9 (Li et al. 2009). We then estimated the average coverage for *GHS-R*, as this was the only gene among those investigated in this work where reads mapped to, using qualimap v.2.2.2-dev (option: *bamqc*; Okonechnikov et al. 2016). We checked for repeat insertions in the area between *IRAK2* and *SEC13*, where *ghrelin* should be situated using RepeatModeler (Flynn et al. 2020) in the garden warbler genome (GCA_014839755.1).

In order to investigate patterns of molecular evolution of the *GHS-R* gene in passerine and non-passerine birds, we downloaded ortholog nucleotide sequences from NCBI (aves, taxid:8782). These were then aligned using MAFFT (Katoh et al. 2009). We only included sequences with a start codon at the beginning and a stop codon at the end of the sequence in the analyses. We further removed any sequences that showed internal stop codons. This resulted in 57 non-passerine and 29 passerine sequences for *GHS-R*. We then carried out molecular evolution analyses using HyPhy (Pond & Muse 2005) via the datamonkey online server (http://www.datamonkey.org/; Weaver et al. 2018) for passerines and non-passerines, respectively. We used the FUBAR (Fast, Unconstrained Bayesian AppRoximation) and the SLAC (Single-Likelihood Ancestor Counting) models (Kosakovsky, Pond & Frost 2005). We next investigated whether the reduced number of sites under purifying selection in passerine birds may result from selection relaxation using RELAX (Wertheim et al. 2015) via HyPhy (Pond & Muse 2005) on the datamonkey online server.

## Acknowledgements

This work was supported by grant N. P 31047-B29 of the Austrian Science Fund (FWF).

## Data Availability

Genomic reads of the three *Sylvia borin* samples can be found on NCBI BioProject PRJNA975133.

## References

Bairlein, F. (1991). Body mass of Garden Warblers (Sylvia borin) on migration: a review of field data. Vogelwarte, 36(1), 48–61.

Battley, P. F., Warnock, N., Tibbitts, T. L., Gill Jr, R. E., Piersma, T., Hassell, C. J., & Riegen, A. C. (2012). Contrasting extreme long-distance migration patterns in bar-tailed godwits Limosa lapponica. Journal of Avian Biology, 43(1), 21–32.

Bushnell, B. (2014). BBMap: a fast, accurate, splice-aware aligner (No. LBNL-7065E). Lawrence Berkeley National Lab.(LBNL), Berkeley, CA (United States).

Dufour, P., Descamps, S., Chantepie, S., Renaud, J., Guéguen, M., Schiffers, K., & Lavergne, S. (2020). Reconstructing the geographic and climatic origins of long-distance bird migrations. Journal of Biogeography, 47(1), 155–166.

Davenport, A. P., Bonner, T. I., Foord, S. M., Harmar, A. J., Neubig, R. R., Pin, J. P., & Kangawa, K. (2005). International Union of Pharmacology. LVI. Ghrelin receptor nomenclature, distribution, and function. Pharmacological reviews, 57(4), 541–546.

Eikenaar, C., Hessler, S., Ballstaedt, E., Schmaljohann, H., & Kaiya, H. (2018). Ghrelin, corticosterone and the resumption of migration from stopover, an automated telemetry study. Physiology & behavior, 194, 450–455.

Flynn, J. M., Hubley, R., Goubert, C., Rosen, J., Clark, A. G., Feschotte, C., & Smit, A. F. (2020). RepeatModeler2 for automated genomic discovery of transposable element families. Proceedings of the National Academy of Sciences, 117(17), 9451–9457.

Friedman-Einat, M., & Seroussi, E. (2019). Avian leptin: bird’s-eye view of the evolution of vertebrate energy-balance control. Trends in Endocrinology & Metabolism, 30(11), 819–832.

Ge, X., Yang, H., Bednarek, M. A., Galon-Tilleman, H., Chen, P., Chen, M., & Kaplan, D. D. (2018). LEAP2 is an endogenous antagonist of the ghrelin receptor. Cell metabolism, 27(2), 461–469.

Goymann, W., Lupi, S., Kaiya, H., Cardinale, M., & Fusani, L. (2017). Ghrelin affects stopover decisions and food intake in a long-distance migrant. Proceedings of the National Academy of Sciences, 114(8), 1946–1951.

Henderson, L. J., Cockcroft, R. C., Kaiya, H., Boswell, T., & Smulders, T. V. (2018). Peripherally injected ghrelin and leptin reduce food hoarding and mass gain in the coal tit (Periparus ater). Proceedings of the Royal Society B: Biological Sciences, 285(1879), 20180417.

Huelsmann, M., Hecker, N., Springer, M. S., Gatesy, J., Sharma, V., & Hiller, M. (2019). Genes lost during the transition from land to water in cetaceans highlight genomic changes associated with aquatic adaptations. Science advances, 5(9), eaaw6671.

Kaiya, H., Furuse, M., Miyazato, M., & Kangawa, K. (2009). Current knowledge of the roles of ghrelin in regulating food intake and energy balance in birds. General and comparative endocrinology, 163(1-2), 33–38.

Kaiya, H., Konno, N., Kangawa, K., Uchiyama, M., & Miyazato, M. (2014). Identification, tissue distribution and functional characterization of the ghrelin receptor in West African lungfish, Protopterus annectens. General and comparative endocrinology, 209, 106–117.

Kaiya, H., Miyazato, M., & Kangawa, K. (2011). Recent advances in the phylogenetic study of ghrelin. Peptides, 32(11), 2155–2174.

Katoh, K., Asimenos, G., & Toh, H. (2009). Multiple alignment of DNA sequences with MAFFT. In Bioinformatics for DNA sequence analysis (pp. 39–64). Humana Press.

Kojima, M., Hamamoto, A., & Sato, T. (2016). Ghrelin O-acyltransferase (GOAT), a specific enzyme that modifies ghrelin with a medium-chain fatty acid. The Journal of Biochemistry, 160(4), 189–194.

Kojima, M., Hosoda, H., Date, Y., Nakazato, M., Matsuo, H., & Kangawa, K. (1999). Ghrelin is a growth-hormone-releasing acylated peptide from stomach. Nature, 402(6762), 656–660.

Kosakovsky Pond, S. L., & Frost, S. D. (2005). Not so different after all: a comparison of methods for detecting amino acid sites under selection. Molecular biology and evolution, 22(5), 1208–1222.

Li, H., & Durbin, R. (2009). Fast and accurate short read alignment with Burrows–Wheeler transform. bioinformatics, 25(14), 1754–1760.

Li, H., Handsaker, B., Wysoker, A., Fennell, T., Ruan, J., Homer, N., & 1000 Genome Project Data Processing Subgroup. (2009). The sequence alignment/map format and SAMtools. bioinformatics, 25(16), 2078–2079.

Lupi, S., Morbey, Y. E., MacDougall-Shackleton, S. A., Kaiya, H., Fusani, L., & Guglielmo, C. G. (2022). Experimental ghrelin administration affects migratory behaviour in a songbird. Hormones and Behavior, 141, 105139.

McGowen, M. R., Tsagkogeorga, G., Williamson, J., Morin, P. A., & Rossiter, A. S. J. (2020). Positive selection and inactivation in the vision and hearing genes of cetaceans. Molecular Biology and Evolution, 37(7), 2069–2083.

Meyer, M., & Kircher, M. (2010). Illumina sequencing library preparation for highly multiplexed target capture and sequencing. Cold Spring Harbor Protocols, 2010(6), pdb-prot5448.

Monroe, J. G., McKay, J. K., Weigel, D., & Flood, P. J. (2021). The population genomics of adaptive loss of function. Heredity, 126(3), 383–395.

Müller, T. D., Nogueiras, R., Andermann, M. L., Andrews, Z. B., Anker, S. D., Argente, J., & Tschöp, M. H. (2015). Ghrelin. Molecular metabolism, 4(6), 437–460.

Murrell, B., Moola, S., Mabona, A., Weighill, T., Sheward, D., Kosakovsky Pond, S. L., & Scheffler, K. (2013). FUBAR: a fast, unconstrained bayesian approximation for inferring selection. Molecular biology and evolution, 30(5), 1196–1205.

Nakazato, M., Murakami, N., Date, Y., Kojima, M., Matsuo, H., Kangawa, K., & Matsukura, S. (2001). A role for ghrelin in the central regulation of feeding. Nature, 409(6817), 194–198.

Okonechnikov, K., Conesa, A., & García-Alcalde, F. (2016). Qualimap 2: advanced multi-sample quality control for high-throughput sequencing data. Bioinformatics, 32(2), 292–294.

Pond, S. L. K., & Muse, S. V. (2005). HyPhy: hypothesis testing using phylogenies. In Statistical methods in molecular evolution (pp. 125–181). Springer, New York, NY.

Prum, R. O., Berv, J. S., Dornburg, A., Field, D. J., Townsend, J. P., Lemmon, E. M., & Lemmon, A. R. (2015). A comprehensive phylogeny of birds (Aves) using targeted next-generation DNA sequencing. Nature, 526(7574), 569–573.

Seim, I., Jeffery, P. L., Herington, A. C., & Chopin, L. K. (2015). Comparative analysis reveals loss of the appetite-regulating peptide hormone ghrelin in falcons. General and Comparative Endocrinology, 216, 98–102.

Selvatti, A. P., Gonzaga, L. P., & de Moraes Russo, C. A. (2015). A Paleogene origin for crown passerines and the diversification of the Oscines in the New World. Molecular phylogenetics and evolution, 88, 1–15.

Tolle, V., Bassant, M. H., Zizzari, P., Poindessous-Jazat, F., Tomasetto, C., Epelbaum, J., & Bluet-Pajot, M. T. (2002). Ultradian rhythmicity of ghrelin secretion in relation with GH, feeding behavior, and sleep-wake patterns in rats. Endocrinology, 143(4), 1353–1361.

Tschöp, M., Smiley, D. L., & Heiman, M. L. (2000). Ghrelin induces adiposity in rodents. Nature, 407(6806), 908–913.

Weaver, S., Shank, S. D., Spielman, S. J., Li, M., Muse, S. V., & Kosakovsky Pond, S. L. (2018). Datamonkey 2.0: a modern web application for characterizing selective and other evolutionary processes. Molecular biology and evolution, 35(3), 773–777.

Wertheim, J. O., Murrell, B., Smith, M. D., Kosakovsky Pond, S. L., & Scheffler, K. (2015). RELAX: detecting relaxed selection in a phylogenetic framework. Molecular biology and evolution, 32(3), 820–832.

