## Supplementary File 1 for "The unexpected loss of the “hunger hormone” ghrelin in true passerines: A game changer in migration physiology"

**Supplement 1. Laboratory methodology – library preparation (modified after Meyer and Kircher 2010)**

**Step 1. Enzymatic fragmentation with dsDNA Fragmentase**

| **reagent** | **volume per sample (μl)** |
| --- | --- |
| 10x buffer | 2 |
| dsDNA Fragmentase | 2 |
| H_2_O | 6 |
| DNA sample | 10 |

Workflow and incubation conditions:

Everything is prepared on ice, thoroughly vortexed, and samples are incubated at 37°C for 40 minutes. Reaction is stopped by addition of 5μl of 0.5M EDTA per sample.

Magnetic clean-up of DNA (after Rohland and Reich, 2012)

Magnetic bead solution (MBS) [10ml]:

| **reagent** | **volume** |
| --- | --- |
| Sera-Mag Speed beads (washed 2x in 500 μl 1x TE buffer) | 100μl |
| PEG-8000 | 1.8g |
| NaCl [5M] | 2.0ml |
| TrisHCl [1M] (pH 8) | 100μl |
| EDTA [100mM] | 100μl |
| Tween20 | 5μl |
| H_2_O | top-up to 10 ml |

Workflow:

- Homogenize MBS by vortexing (avoid foam formation)
- Add 2 sample-volumes of MBS to the samples
- Vortex and incubate at RT for 5 min
- Place on magnet and wait until the liquid is clear and the beads are attached to the tube wall
- Discard supernatant (while the samples are still on the magnet)
- Leave samples on the magnet and add 200 μl EtOH [70%] and pipet up and down 3 times
- Discard supernatant and repeat the wash step with 120-200 μl EtOH [70%](leave the samples all the time on the magnet)
- Discard supernatant and make sure that there is no EtOH remaining
- Air-dry the samples for 5-10 minutes (don’t overdry)
- Add 34 μl [this volume depends on your desired end-volume] EBT (or 1x TE or AE-buffer [Qiagen]) and mix by vortexing
- Incubate for 5 min at 37°C or overnight in the fridge and put back on the magnet
- Collect 32 μl [this volume can be changed as well] of the clear eluted sample and place in a new tube

**Step 2. Blunt-end repair**

| **reagent** | **volume per sample (μl)** |
| --- | --- |
| T4 DNA polymerase [3U/μl] | 0.4 |
| T4 PNK [10U/μl] | 1 |
| dNTP’s 25mM | 0.4 |
| T4 DNA ligase buffer [10x] | 4 |
| reaction enhancer | 2.2 |
| DNA template | 32 |

Workflow and PCR conditions:

Mastermix is prepared on ice.

| **Temperature (°C)** | **time** |
| --- | --- |
| 20 | 30 min |
| 65 | 30 min |
| 4-12 | ∞ |

**Step 3. Adapter ligation**

Use double stranded mp5 adpaters:

| **reagent** | **volume per sample (μl)** |
| --- | --- |
| T4 DNA ligase buffer [10x] | 1 |
| PEG-4000 [50%] | 6 |
| T4 DNA ligase [400U/μl] | 1 |
| mP5 adapter mix | 2 |
| DNA template | 40 |

Workflow and PCR conditions:

Mastermix is prepared on ice.

| **Temperature (°C)** | **time** |
| --- | --- |
| 20 | 30 min |
| 65 | 10 min |
| 4-12 | ∞ |

**Step 4. Size selective purification with ProNex chemistry (Promega)**

Follow protocol 6A from the kit instructions using 1.2x Pronex Chemistry

**Step 5. Indexing PCR**

| **reagent** | **volume per sample (μl)** |
| --- | --- |
| All Taq Mix (4x) | 5 |
| P5 index | 0.5 |
| P7 index | 0.5 |
| H_2_O | 4 |
| DNA template | 10 |

PCR conditions:

| **Temperature (°C)** | **time** |
| --- | --- |
| 95 | 2 min |
| 95 | 5 sec |
| 56 | 15 sec |
| 72 | 10 sec |
| 4-12 | ∞ |

- Repeat 20x

**Step 6. Pooling of samples for library**

We pooled all samples after indexing and divided the pool into three parts for a gel extraction in which we cut out pieces with an approximate median fragment length of 450bp. We performed the gel extraction with a kit (Analytik Jena) and followed the manufacturer’s protocol. In parallel, we also performed a gel extraction after Sun et al., 2012 and used the in-house magnetic bead clean-up protocol as described above. For the actual sequencing library pool we used 60μl of product from each gel extraction method and calculated the molarity of the pool with the Illumina molarity calculator.

**References**

Meyer, M., & Kircher, M. (2010). Illumina sequencing library preparation for highly multiplexed target capture and sequencing. Cold Spring Harbor Protocols, 2010(6), pdb-prot5448.

Rohland, N., & Reich, D. (2012). Cost-effective, high-throughput DNA sequencing libraries for multiplexed target capture. *Genome research*, *22*(5), 939-946.
